# *Correspondence: Spontaneous secondary mutations confound analysis of the essential two component system WalKR in* Staphylococcus aureus

**DOI:** 10.1101/058842

**Authors:** Ian R. Monk, Benjamin P. Howden, Torsten Seemann, Timothy P. Stinear

## Abstract

Ji *et al.*,^1^ recently detailed the structure of the extracytoplasmic Per-Arnt-Sim (PAS) domain of WalK (WalK^EC-PAS^), the sensor kinase of the essential two-component system WalKR in *Staphylococcus aureus*. The authors made two independent walK mutants in *S. aureus*, each with a single amino acid alteration in WalK^EC-PAS^. They postulated from comparative structural analysis and primary sequence comparisons that these residues might be important for extra cellular signal transduction. We have also been exploring the function of WalKR and were surprised by the striking phenotypic impact of a single amino acid substitutions in the WalK sensor, which were contrary to our own unpublished observations.

The authors subjected their WalK^EC-PAS^ mutants (WalK^D119A^ and WalK^Vl49A^) to a series of phenotypic screens to probe the function of this domain. Compared to the parental methicillin sensitive *S. aureus* strain Newman ^2^, the mutants showed dramatic phenotype changes that included reduced susceptibility to lysostaphin exposure, loss of haemolysis on sheep blood agar, reduced biofilm formation and virulence in a mouse infection model. RNA-seq comparisons of mutants to wild type showed substantial transcriptional changes, with 285 (111+174) differentially expressed genes between the two WalK^EC-PAS^ mutants and wild type. They also used structure-based virtual screening to identify 2, 4-dihydroxybenzophenone (DHBP) as a small molecule predicted to interact with the WalK^EC-PAS^ domain. As DHBP stimulated lysostaphin induced lysis and biofilm formation in strain Newman they postulated it as an activator of WalKR. They then compared transcriptional responses of Newman against DHBP exposure and with their D119A mutant. The authors reported that there were 41 genes differentially and inversely expressed in the WalK^D119A^ mutant and DHBP-treated cells, and concluded that this supported a role for DHBP in activating WalKR. No direct evidence to support activation of WalKR through enhanced phosphotransfer was presented^1^.

To investigate further, we were provided with *S. aureus* Newman wild type, WalK^D119A^ and WalK^Vl49A^ by the senior author, Chuan He, University of Chicago (UoC)^1^. We first sequenced the genomes of University of Melbourne (UoM) and UoC Newman strains and compared their sequences to the published reference, with the two strains differing by only one synonymous mutation in *sbnF* (NWMN_0065), showing that the strains were essentially identical. Then, using D119A allelic exchange, we recreated the WalK mutation in the Newman wild type and the USA300 lineage strain NRS384^3^. We confirmed by genome sequencing that only the AT→CG substitution in WalK^EC-PAS^ was introduced in both strains at position 25994 (Accession: PRJEB14381). We then tested these mutants (UoM Newman WalK^D119A^ and NRS384 WalK^D119A^) for some of the key phenotype changes observed by Ji *et al.*,^1^. However, in contrast to Ji *et al.*, ^1^ we observed that UoM WalK^D119A^ and NRS384 WalK^D119A^ were fully hemolytic (Fig. 1A) and exhibited identical growth curve kinetics as the parental strains (optical density and CFU) (Fig. 1B). Concurrent ni 1 QA screening of UoC Newman WalK confirmed it was non-hemolytic (Fig. 1A). The mutant also grew to an increased OD_600_, as reported (Fig. 1B) *^1^,* although CFU counts were identical to wild type, suggesting that the increase in OD is not due to increased growth. Additionally, UoC Newman WalK^D119A^ consistently exhibited larger colonies than Newman or UoM WalK^D119A^. We next measured the sensitivity of the strains to lysostaphin by cell viability (Fig. 1C). We observed that the UoC WalK^D119A^ mutant was significantly more sensitive (not resistant) to lysostaphin than wild type (3-log_10_ reduction vs Newman), whereas the UoM WalK^D119A^ mutant showed no change. Interestingly, the WalK^D119A^ mutation in NRS384 caused an increase in lysostaphin sensitivity, suggesting the mutation contributes to WalKR activation rather than repression, as proposed by Ji *et al.*,^1,4^.

**Fig. 1.**
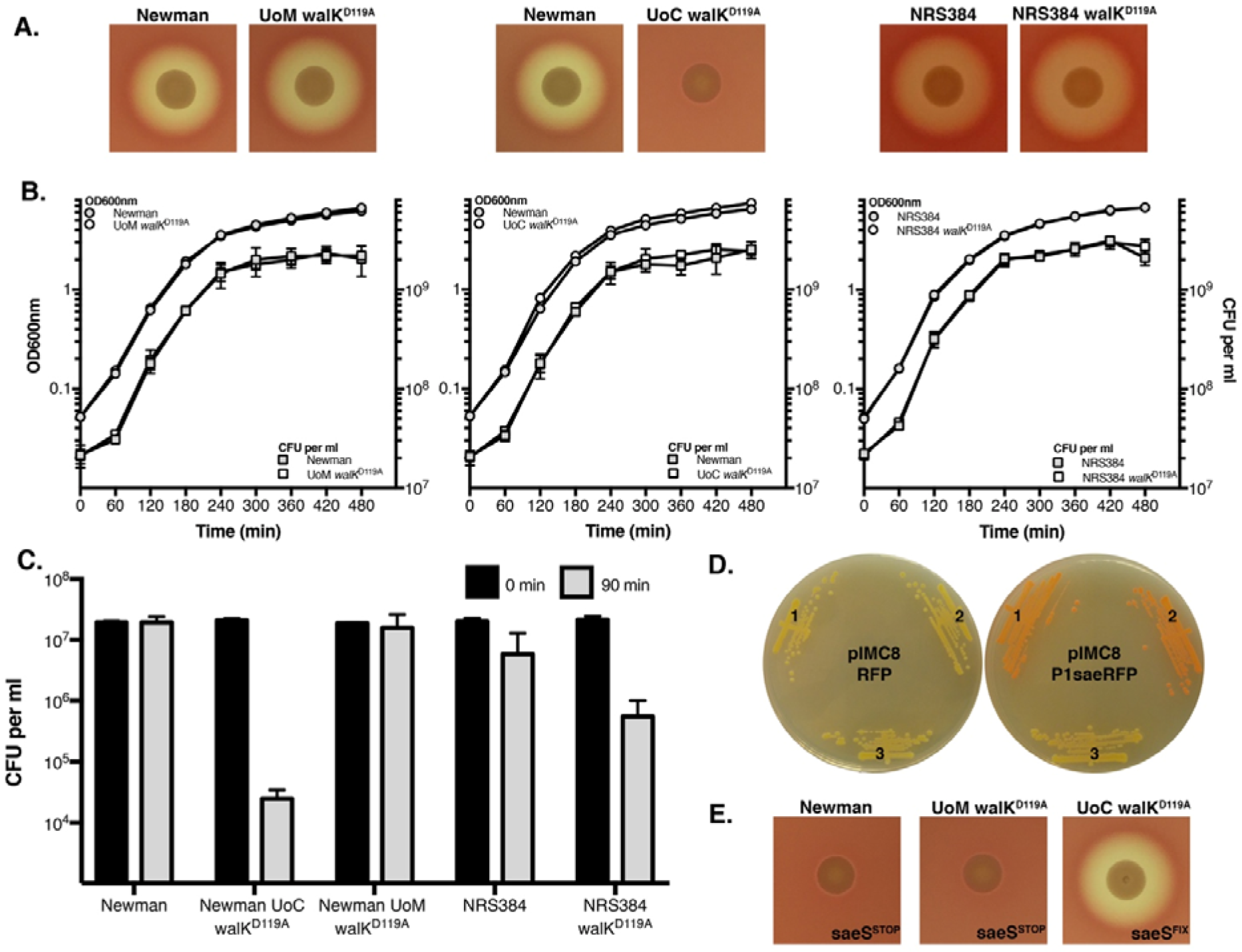
Phenotypic screening of *walK^EC-PAS^* 266 mutants. (A) No impact on haemolysis was observed for the WalK^D119A^ mutation in either Newman or NRS384 on sheep blood agar. (B) Growth kinetics were identical for the newly created WalK^D119A^ 268 mutants in TSB at 37°C with aeration (200 rpm) when compared to the parental strain. Optical density (O) and colony forming units were enumerated. (C) In a lysostaphin growth sensitivity assay the UoC WalK^D119A^ exhibited a loss of viability, while the parent or UoM WalK^D119A^ did not. The WalK^D119A^ mutation in the NRS384 background enhanced sensitivity to lysostaphin when compared to the parental strain. Error bars depict the standard deviation of the mean from three independent experiments. (D) P1 sae promoter activity with a DsRED reporter. Left: no promoter. Right: P1 sae driving RFP expression; (1) Newman (2) UoM WalK^D119A^ (3) UoM WalK^D119A^. No expression of *sae* was observed in UoC WalK^D119A^. (E) By allelic exchange the UoC *saeS* mutation was introduced into Newman or UoM WalK^D119A^ (saeSSTOP277) abolished haemolysis and the introduction of the wild type *saeS* gene into UoC WalK^D119A^ (*saeS*^FIX^) restored haemolysis.

To resolve the discrepancy between the published results ^1^ and our observations, we subjected UoC WalK^D119A^ and UoC WalK^V149^ to whole genome sequencing and identified all DNA differences (Accession: PRJEB14381). Relative to Newman - and in addition to their expected *walK^EC-PAS^* changes - both UoC mutants D119A and V149A had acquired six additional mutations (Table 1), most notably two independent loss-of-function mutations in *saeRS,* a major two-component regulator that controls expression of many genes involved in virulence and biofilm formation ^5–8^.

**Table 1:**
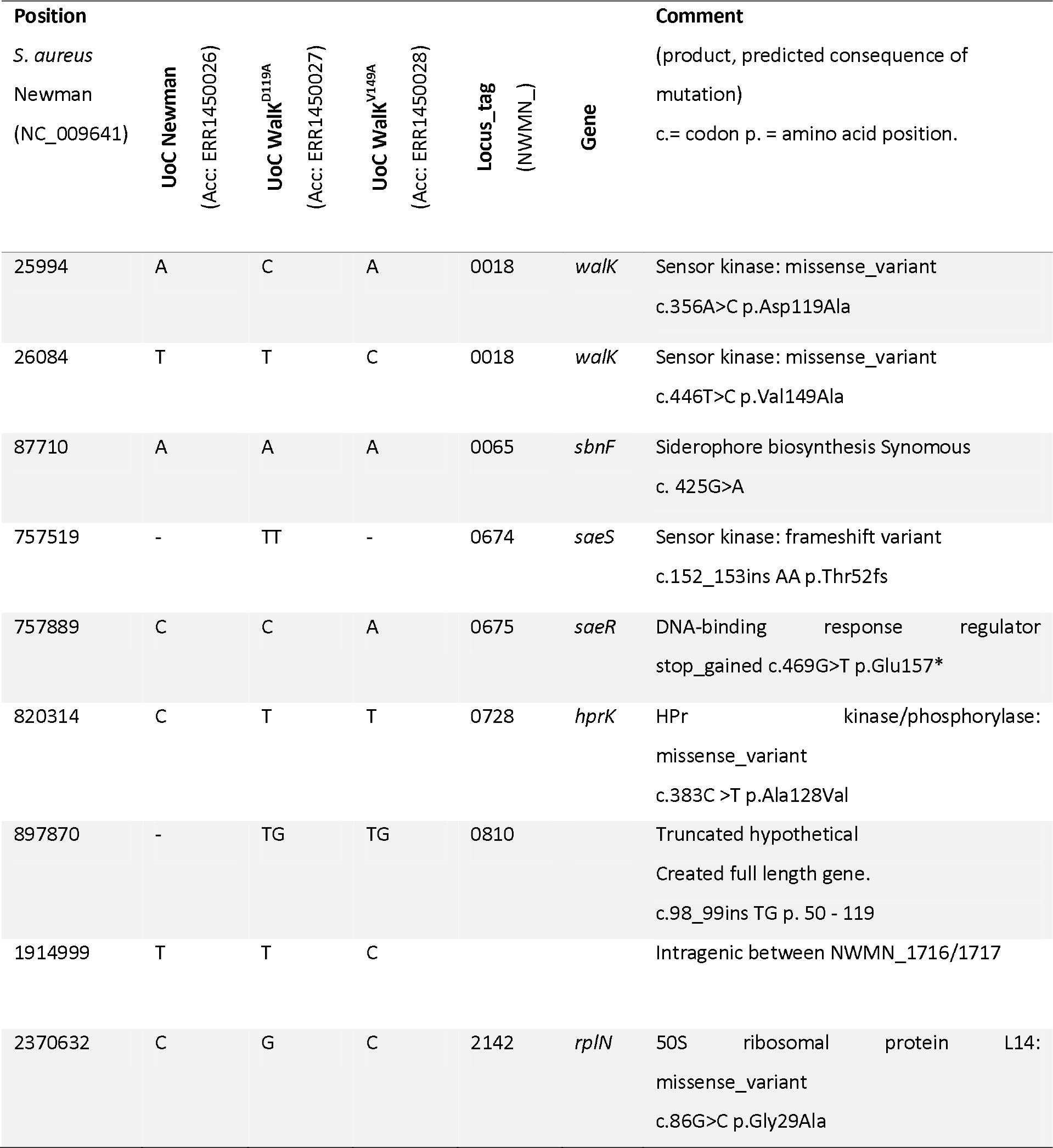
Whole genome sequencing to identify the introduction of unintended polymorphisms within the *S. aureus* Newman WalK mutants.

The UoC WalK^D119A^ had a TT insertion at chromosome position 757519 that introduced a frameshift to *saeS*. The UoC WalK^vl49A^ had a G→T substitution at chromosome position 757889 that introduced a premature stop codon to *saeR*. It is these secondary mutations in *saeRS*, rather than the targeted mutations in *walK^EC-PAS^*, that likely explain the phenotypes observed by Ji *et al.*, (reduced biofilm, loss of haemolysis, reduced virulence)^1^. To confirm the predicted functional consequences of the *saeRS* mutations, we used a P1 Sae red fluorescence reporter plasmid ^5^. No fluorescence activity was detected in the UoC WalK^D119A^ strain containing the P1 Sae reporter, consistent with the predicted truncated histidine kinase preventing phosphorylation of SaeR. While high level expression of P1 Sae from Newman and UoM WalK^D119A^ was observed leading to red colonies (Fig. 1D) and high level fluorescence expression (data not shown). We then recreated the mutated *saeS* allele from UoC WalK^D119A^ in both wild type Newman and UoM WalK^D119A^ (Fig. 1E). The mutation abolished haemolysis on sheep blood agar. We then repaired the *saeS* mutation in UoC WalK^D119A^ and observed restoration of wild type haemolysis (Fig. 1E). These results categorically show that the UoC WalK^D119A^ strain is an *sae* mutant with the majority of the phenotypic changes reported in this strain (including the RNAseq changes discussed below) likely associated with this mutation rather than the WalK^D119A^ mutation. These unintended secondary mutations in a major *S. aureus* regulatory locus preclude analysis of the role of WalK^EC-PAS^ domain in WalKR signal transduction.

There is precedence for this specific phenomenon. Sun *et al.*, showed in detail that elevated temperature and antibiotic selection used during the *S. aureus* mutagenesis process can aid in the selection of *saeRS* mutations ^5^. How Ji et al., managed to complement the mutations (D119A and V149A) by phage integrase plasmid expression (pCL55) of wild type *walKR* remains to be explained^1^. We have been unable thus far to obtain the complemented mutants from the authors for further analysis.

We also observed that UoC WalK^D119A^ and UoC WalK^Vl49A^ exhibited larger colonies compared to wild type, a phenotypic difference not discussed by Ji *et al.*^1^. This change in both mutants might be explained by the C→T substitution observed at 820314, leading to an A128V change in HprK, a serine kinase known to be involved in catabolite repression and associated with a spreading colony phenotype^9^.

We next re-examined the authors’ RNAseq data (Accession: GSE75731) obtained from biological duplicate experiments of *S. aureus* Newman wild type, WalK^D119A^, WalK^Vl49A^ and Newman with 50µM DHBP treatment. Using *Kallisto*^10^ and the *S. aureus* Newman reference genome (Accession: NC_009641) we quantified transcript abundance for each of the four conditions. The output of *Kallisto* was analysed and visualized using Degust ^11^, the latter an interactive website to explore the analysis ^12^. Multi-dimensional scaling plots showed good consistency between the biological replicates, with Newman wild type, Newman-treated-with-DHBP and the two mutants each displayed clustering indicative of distinct transcriptional profiles (Fig. 2a). Applying the same threshold cutoffs (2-fold change) and including a false discovery rate (FDR) of <0.01, we then replicated the intersection analysis of Ji *et al.*, but found only nine genes (not the reported 41) that were down regulated in WalK^D119A^ and up regulated in Newman treated with DHBP (Fig. 1C). Among the nine genes, only five overlapped with the 41 reported by Ji *et al.*,^1^ and none of these included autolysins previously linked to the WalKR regulon (Table 2)^13,14^. In fact, five of the nine genes listed have all previously been reported to be under SaeRS control (Table 2). We repeated our lysostaphin assay with and without the addition of 75µM DHBP, however we did not observe an impact on lysostaphin induced cell viability which is in direct contrast to the results of Ji *et al.*^1^ where they observed increased loss of turbidity in Newman pre-treated with DHBP upon lysostaphin treatment.

**Fig. 2.**
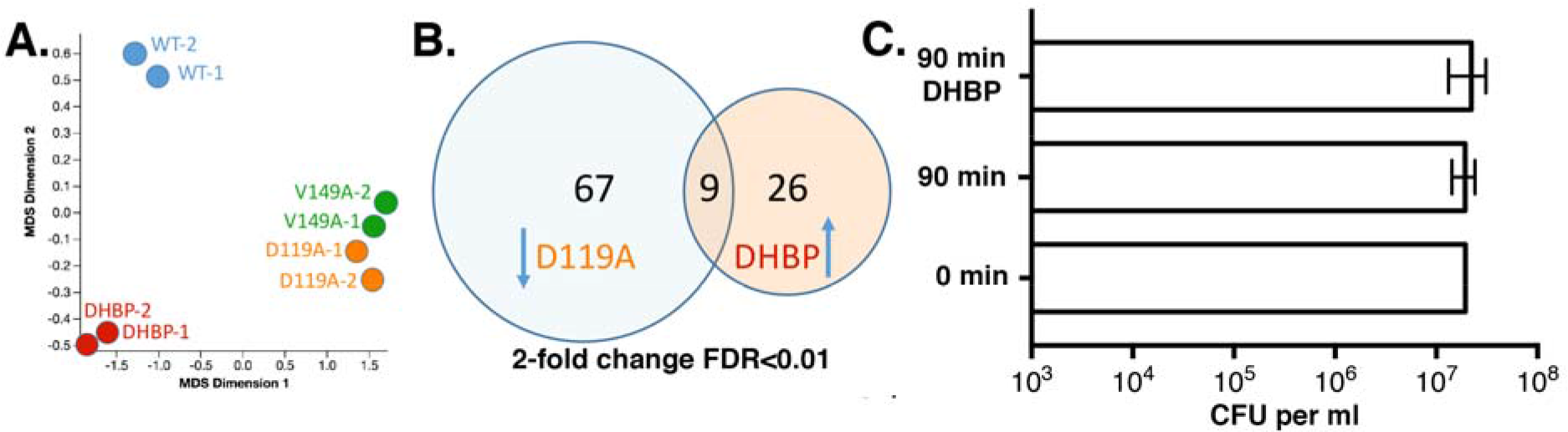
Comparisons of *S. aureus walK*^*EC*-PAS^ mutants. Multidimensional scaling plot based on published RNAseq data from Ji et al ^1^, showing similar transcriptional profiles for the *walK*^*EC*-PAS^ mutants but distinct from wild type treated with 2, 4-dihydroxybenzophenone (DHBP). (B) Venn diagram showing nine CDS that are down regulated in the D119A mutant and upregulated on exposure to DHBP. (C) No enhanced impact on lysostaphin induced lysis in strain Newman was observed when grown in the presence of 75 µM DHBP.

**Table 2:** List of the nine CDS shown in Fig. 1B that were down regulated in D119A and up-regulated on exposure to DHBP (both treatments compared to *S. aureus* Newman wild type).

| No. | Product | Locus_tag<br>NWMN_ | Log<sub>2</sub>FC<br>WT vs<br>D119A | Log<sub>2</sub>FC<sup>#</sup><br>WT vs<br>DHBP |
| --- | --- | --- | --- | --- |
| **1<sup>§</sup>** | Staphylocoagulase precursor (*coa*) | 0166 | -2.92 | 2.40* |
| **2** | Superantigen-like protein 7 (*ssl7*) | 0394 | -1.64 | 2.46* |
| **3** | Hypothetical protein | 0401 | -1.94 | 2.56 |
| 4 | Sodium-dependent symporter protein | 0423 | -1.18 | 1.47 |
| **5** | Secreted VWF-binding protein precursor | 0757 | -1.88 | 2.40* |
| 6 | Cytochrome D ubiquinol oxidase, subunit I | 0952 | -2.05 | 1.32 |
| 7 | Cytochrome D ubiquinol oxidase, subunit II | 0953 | -1.88 | 1.39 |
| **8** | Fibronectin binding protein B (*fnbB*) | 2397 | -4.02 | 1.53* |
| 9 | Immunodominant antigen A (*isaA*) | 2469 | -1.25 | 1.45* |

Notes: ^#^FDR <0.01; ^§^Numbers in bold typeface indicate previously reported as SaeRS regulated ^5,6^. *Asterisk indicates also reported as upregulated by Ji *et al.*^1^.

We also noted that only 52 genes were differentially expressed upon exposure to DHBP, in comparison to the reported 145 genes ^1^. We think this discrepancy and the preceding difference were due to the absence of any filter for false discovery rate (FDR) applied by Ji *et al.*,^1^. For example, our analysis of their RNA-seq data without a FDR threshold resulted in 172 differentially expressed genes in Newman treated with DHBP compared to Newman alone ^12^. Transcriptome data without statistical significance thresholds are not meaningful^15^.

We also mapped the authors’ RNAseq reads for WalK^D119A^ and WalK^Vl49A^ (Accession: GSE75731) to the *S. aureus* Newman reference (Accession: NC_009641) and readily detected the same saeRS mutations we observed from our independent sequencing of these mutants.

## Conclusion

Our analyses highlight two major issues with the study of Ji *et al.*^1^ Firstly, the presence of unintended *saeRS* mutations in their *walK*^EC-PAS^ mutants invalidate their conclusions with respect to role of the WalK extracytoplasmic domain in controlling WalKR function. We observed opposing results; with increased sensitivity to lysostaphin of the clean D119A mutant, a phenotype previously linked with enhanced activity of the system ^4^. Secondly, the RNAseq data appears to have been inadequately analysed, with no filtering for false discovery. Application of an appropriate threshold (p<0.01) to their data substantially changes the lists of differentially expressed genes. The authors’ use the overlap between the D119A mutant and DHBP treatment to ‘prove’ that DHBP is signaling through WalK^EC-PAS^, but this conclusion is not supported by the data.

The discovery of small molecule inhibitors of WalKR function would represent a major advance in the fight against multidrug resistant *S. aureus,* and Ji *et al.*,^1^ have shown a potential approach through establishing the structure of WalK^EC-PAS^. Unfortunately, based on the data presented, the authors’ principal conclusions regarding WalK^EC-PAS^ domain function are not supported. This study is another example of the pitfalls associated with allelic exchange in *S. aureus* and the rigor that must applied to mutation validation ^5^. Whole genome sequencing is now so affordable that it can be readily used to verify targeted mutants and the complemented strains. In our own WalKR research we have observed a propensity for mutations introduced into this locus to yield secondary compensatory events ^13,16^. These secondary changes can confound analysis of this essential two-component system, and highlight the extreme care needed when manipulating this locus and then attributing specific phenotypes to specific mutational changes.

## Methods

### Bacterial strains, primers, plasmids and growth conditions

The *S. aureus* UoM Newman was obtained from Prof. Tim Foster (Trinity College Dublin), NRS384 was obtained from BEI resources (www.beiresources.org). *S. aureus* was routinely grown in Tryptic Soy Broth (TSB-Oxoid) at 37 °C with aeration at 200 rpm. Primers were purchased from IDT (www.idtdna.com) with primer sequences detailed in Table 3. Restriction enzymes, Phusion DNA polymerase and T4 DNA ligase were purchased from New England Biolabs. Genomic DNA was isolated from 1 ml of an overnight culture (DNeasy blood and tissue kit - Qiagen) pretreated with 100µg of lysostaphin (Sigma). DHBP was purchased from Sigma (cat no. 126217-lOOg).

**Table 3:**
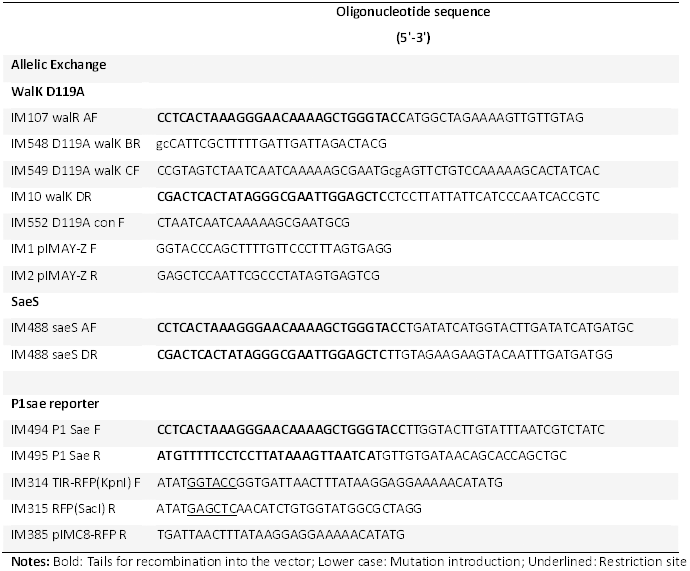
Oligonucleotides used in the study.

**Notes:** Bold: Tails for recombination into the vector; Lower case: Mutation introduction; Underlined: Restriction site

### Lysostaphin sensitivity assay

Overnight cultures of *S. aureus* were diluted 1:100 in fresh, pre-warmed TSB in the presence of 0.2µg/ml of lysostaphin (AMBI) with or without 75µM DHBP (100 mM stock in methanol). Broths were incubated statically at 37°C. Colony forming units were determined by spot plate dilution on Brain heart infusion agar at 0 and 90 min. Limit of detection for the assay was 10^3^ CFU/ml.

### Construction of plMC8-RFP and SLiCE cloning

The *S. aureus* codon optimized DsRED red fluorescent protein and upstream TIR sequence from pRFP-F ^17^ was PCR amplified with primers IM314/IM315. The product was digested with Kpnl/Sacl and cloned into the complementary digested pIMC8 (non-temperature sensitive version of pIMC5 ^18^, creating pIMC8-RFP. To clone into pIMAY-Z ^2^ and plMC8-RFP primers were tailed with 30 nt of complementary sequence to the plasmid. Amplimers were inserted with seamless ligation cloning extract (SLiCE) ^19^ into the vector (pIMAY-Z: *walRK^D119A^, sae^STOP^, sae^FLX^*; plMC8-RFP: P1 sae). Either vector was linearized with KpnI, gel extracted and PCR amplified with primers IM1/IM2 (pIMAY-Z) or IM1/IM385 (pIMC8-RFP). Both amplimers (vector and insert) were combined in a 10µl reaction containing 1×T4 ligase buffer, with 1µl of SLiCE extract. The reaction was incubated at 37 for 1h and then transformed into *Escherichia coli* strain IM08B ^2^, with selection on Luria agar plates containing chloramphenicol 10 µg/ml. Plasmids were extracted and directly transformed by electroporation into the target *S. aureus* strain ^2^.

### Production of SLiCE extract

The SLiCE was isolated from DY380 grown in 50 ml 2xYT at 30°C after a 1:100 dilution of the overnight culture and a 42°C heat shock (50 ml of 42°C 2xYT added) for 25 min once the culture reached OD600=2.5. Cells were processed as described by Zhang et al.,^19^ with the pellet lysed in 500 ul of CelLytic B cell lysis reagent (C87040-10ml-Sigma).

### Whole genome sequencing and data analysis

Whole genome sequencing was performed using the Illumina NextSeq (2x150 bp chemistry), with library preparation using Nextera XT (Illumina). Resulting reads were mapped to the *S. aureus* Newman reference (Accession:NC_009641) using *Snippy* v3.1 (https://github.com/tseemann/snippy). Note that 89 substitutions, 20 deletions and 25 insertions were shared between UoC and UoM Newman strains compared to the NC_009641 reference sequence, representing likely sequencing errors in the 2008 published reference^2^.

